# High-confidence 3D template matching for cryo-electron tomography

**DOI:** 10.1101/2023.09.05.556310

**Authors:** Sergio Cruz-León, Tomáš Majtner, Patrick C. Hoffmann, Jan Philipp Kreysing, Maarten W Tuijtel, Stefan L Schaefer, Katharina Geißler, Martin Beck, Beata Turoňová, Gerhard Hummer

## Abstract

Cryo-electron tomography (CryoET) resolves individual macromolecules inside living cells. However, the complex composition and high density of cells challenge the faithful identification of features in tomograms. Here, we capitalize on recent advances in electron tomography and demonstrate that 3D template matching (TM) localizes a wide range of structures inside crowded eukaryotic cells with confidence 10 to 100-fold above the noise level. We establish a TM pipeline with systematically tuned parameters for automated, objective and comprehensive feature identification. High-fidelity and high-confidence localizations of nuclear pore complexes, vaults, ribosomes, proteasomes, lipid membranes and microtubules, and individual subunits, demonstrate that TM is generic. We resolve ∼100-kDa proteins, connect the functional states of complexes to their cellular localization, and capture vaults carrying ribosomal cargo *in situ*. By capturing individual molecular events inside living cells with defined statistical confidence, high-confidence TM greatly speeds up the CryoET workflow and sets the stage for visual proteomics.

## Introduction

Cryo-electron tomography (CryoET) images the cellular environment *in situ* without labels and with fully preserved context^1,2^. Recent advances in hardware and acquisition techniques have enabled CryoET to routinely image, with high throughput, cell volumes in their native state and obtain structures of abundant macromolecular complexes with near molecular resolution^3–5^. However, lacking a uniform established method, the localization of particles in the tomograms remains highly customized, specific to each target, at best semi-automatic, and relying on strong manual input (such as definition of geometric surface for large pleomorphic assemblies)^6–10^ or extensive, often manual corrections for the false positives in an initial automated assignment^3,4,11^. The confident identification of a sufficient number of particles for a challenging target such as the nuclear pore complexes (NPC) can thus take months or years of manual annotation of literally hundreds of tomograms^12,13^. Lacking an automated, general and reliable localization method, we also have not yet realized the promise of visual proteomics^14–17^ to build molecularly detailed representations of complex cellular landscapes from CryoET data.

Reliable assignment of molecular identities in tomograms is challenging due to both the biological context and the specifics of CryoET processing. Cells are crowded environments, and the proteins within them are structurally heterogeneous and vary widely in size and abundance. The physical limitations of the acquisition procedure further complicate particle localization^18–20^. In CryoET, the electron dose is limited to prevent sample radiation damage, which results in a low signal-to-noise ratio in the acquired tilt series. The maximum sample tilt of about ±60 degrees results in incomplete angular sampling known as the “missing wedge” problem in the 3D reconstruction. In addition, the electron micrographs are conventionally captured out of focus. To recover the high-resolution information, it is thus necessary to accurately determine the defocus and correct the contrast transfer function (CTF)^20^. Visual proteomics needs to overcome these challenges for the reliable assignment of molecular identities to noisy three-dimensional (3D) images of highly complex cellular volumes.

Manual tomogram annotation is still widely used despite being labor intensive, intrinsically subjective and incomplete^21^. Template-based computational approaches^22^ use known objects (templates) and compare them with the data by calculating a similarity metric (usually a constrained cross-correlation) ^15,16,23–25^. In contrast, template-free methods iterate to cluster particles and determine patterns without imposing any structure^26,27^. However, their accuracy and efficiency need improvement^22^. Deep learning algorithms, including classification and semantic segmentation, have been applied to CryoET^22,28–30^. Recently, implementations such as DeepFinder^29^, DeePiCt^30^ and TomoTwin^31^ have shown promising results in segmenting tomograms and identifying the positions of common macromolecular complexes.

However, these methods require extensive annotations for training and are less effective in detecting low-abundance particles^22^, so far limiting their use to detect ribosomes and similarly sized particles. Furthermore, they determine only positions and further processing is needed to determine particle orientations.

Template matching (TM)^15,16,23–25^ is typically used with low-resolution templates of the macromolecular complex of interest on down-sampled tomograms to reduce computational cost and avoid template bias. Large numbers of false positive hits are removed either manually, thereby lowering the objectivity of the approach, or through a multistep classification procedure, which is computationally expensive and can fail if the number of particles is small. In addition, the data down-sampling limits the ability to localize smaller or weak-signal particles^32^. In theory, the ability of TM to localize the particles with high confidence should be connected to the quality of the template and how well it resembles the actual data. However, in practice, it has not been objectively shown how TM depends on the type of template and parameters such as voxel size, masks, resolution filtering, and the number of orientations.

In this study, we establish a high-confidence TM pipeline and combine it with CryoET imaging for visual proteomics of eukaryotic cells. We show that the performance of TM not only depends on the size, origin and shape of the template, but also on angular increment in orientational sampling, and on tomogram voxel size (magnification), filtering and resolution. We demonstrate the power of optimized TM to localize nuclear pore complexes (NPCs), vault proteins, ribosomes, proteasomes, microtubules, and lipid membranes, inside a single dataset. We establish that TM can identify low-abundance and low-density complexes with high fidelity, as exemplified by the identification of ribosome-loaded vaults. We show that TM quantitatively captures the diversity of eukaryotic ribosomes in different functional states. TM localizes individual subunits of the NPC, microtubule protofilaments, and the large and small ribosomal subunits. We provide recommendations for users to optimally set template depended search parameters and a parameter estimation software tool.

## Results

### High-confidence template matching for *in situ* macromolecule localization

We comprehensively tested our TM pipeline on tomograms of *D. discoideum* (EMD-XXXX) obtained from lamellae milled with cryo-focused ion beam microscopes^4^ (see Fig. 1 for the workflow and Methods for details on data acquisition). Starting from a library of the best available templates for a series of candidate features, we performed TM of each template in a tomogram independently and assigned particle identities to the points with high constrained cross-correlation (CC). The locations and orientations of the assigned peaks permit the visualization and analysis of the spatial interactions of the features. We used a total of 18 templates (Table 1) at different voxel sizes and with multiple search parameters including the number of orientations and filters (see Methods for details). Templates in the library were obtained from different sources including subtomogram averaging (STA), homology modelling, the protein data bank (PDB)^33^, the electron microscopy data bank (EMDB)^34^, and molecular dynamics simulations (see Methods for details).

**Fig. 1:**
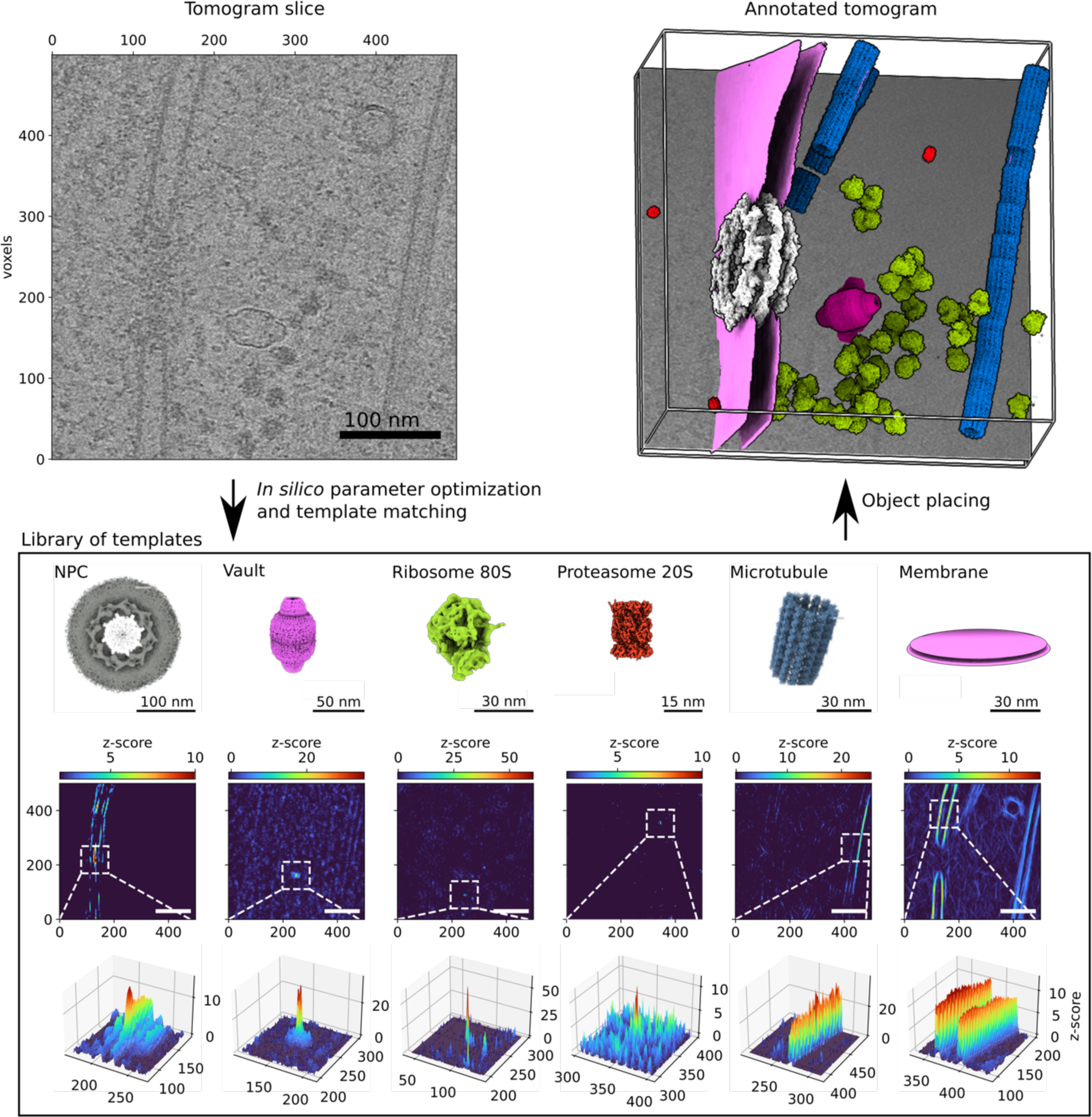
Template matching for visual proteomics. A tomogram (slice shown in top left) is cross-correlated independently with each template in the library (bottom) to identify points with high constrained cross-correlation values (zoom-ins with CC z-scores at bottom). From the z-score maps, three-dimensional localization maps (top right) are generated for visualization^58^ and analysis of the spatial interactions of proteins and their complexes.

**Table 1:** Tested cases for template matching.

| Template | method | reference | angle sampling<br>(deg) | symmetry | size<br>(voxels<sup>3</sup>) | Tilt series | magnification<br>(Å/voxel) |
| --- | --- | --- | --- | --- | --- | --- | --- |
| NPC (*D. discoideum*) | STA | EMD-XXX | 5 | c8 | 240 | 126, 89 | 8.70 |
| NPC subunit (*D. discoideum*) | STA | EMD-XXX | 10 | c1 | 124 | 89, 126 | 8.70 |
| NPC subunit (*Homo sapiens*) | PDB | PDB-id: 7R5J | 10 | c1 | 124 | 126 | 8.70 |
| Modelled vault protein (*D. discoideum*) | Swiss-model | This work | 10 | c39 | 100 | 126, 90 | 8.70 |
| Modelled half vault (*D. discoideum*) | Swiss-model | This work | 10 | c39 | 100 | 126, 90 | 8.70 |
| STA Vault protein (*D. discoideum*) | STA | This work | 10 | c39 | 100 | 126, 90 | 8.70 |
| Ribosome 80S (*D. discoideum*) | STA | EMD-15810 | 30, 20, 10, 5 | c1 | 48, 100 | 126, 109, 89 | 8.70, 4.35 |
| SRS 40S (*D. discoideum*) | STA | This work | 10 | c1 | 100 | 89 | 4.35 |
| LRS 60S (*D. discoideum*) | STA | This work | 10 | c1 | 100 | 89 | 4.35 |
| Unrotated ribosome (*D. discoideum*) | STA | EMD-15812 | 10 | c1 | 100 | 109 | 4.35 |
| Rotated ribosome (*D. discoideum*) | STA | EMD-15815 | 10 | c1 | 100 | 109 | 4.35 |
| Proteasome 20S (*Homo sapiens*) | PDB | PDB-id: 6RGQ, EMD-4877 | 10 | c1 | 38, 76 | 27 | 8.70, 4.35 |
| Microtubule (*Homo sapiens*) | PDB | PDB-id: 3JAR, EMD-6351 | 10 | c13 | 52 | 195 | 8.70 |
| Protofilament (*Homo sapiens*) | PDB | PDB-id: 3JAR, EMD-6351 | 10 | c1 | 52 | 195 | 8.70 |
| Membrane (small STA) | STA | This work | 20, 10, 2 | c360 | 50 | 126 | 8.70 |
| Membrane (large STA) | STA | This work | 20, 10, 2 | c360 | 50 | 126 | 8.70 |
| Membrane from MD (atomic model) | MD simulation | This work | 20, 10, 2 | c360 | 50 | 126 | 8.70 |
| Half proteasome (*Homo sapiens*) | PDB | PDB-id: 6RGQ, EMD-4877 | 10 | c1 | 42, 84 | 027 | 4.35, 2.176 |

We used the STOPGAP^35^ software framework to calculate the actual cross-correlation between templates and tomograms, which maximizes the cross-correlation of the template according to its orientation and positions, and already takes missing wedge, angular tilt step, defocus, and electron dose into account (see Methods for details). For each template, with optimized search parameters (see next section), peaks several standard deviations above noise appear in the z-score map. High-confidence peaks correspond to the position where the center of the template is placed to best reproduce the data from the tomograms.

Fig. 1 summarizes the TM procedure. We used a library that includes templates for the NPC, the 80S ribosome, and the nuclear envelope obtained by STA from tomograms of *D. discoideum*. For the proteasome^36^ and microtubule^37^, we used the human homologous structures previously reported (PDB-id: 6rgq, PDB-id: 3jar, respectively). For the vault, we created a density map starting from an atomic model generated by homology modeling. With each of the templates we performed TM, initially at 4-binned data with a voxel size of 8.704 Å and then also at higher resolution (2-binned 4.352 Å/voxel and unbinned 2.176 Å/voxel). By progressing hierarchically to higher resolution, we aimed to capitalize on the high signal content of the data collected with latest generation hardware.

We transformed the cross-correlation volumes into z-score maps. In this representation, a peak at the center of the NPC is typically ∼10 standard deviations (𝜎) above the map noise, while the vault and the ribosome have peaks with CC values of ∼30 𝜎 and ∼40 𝜎, respectively (Fig. 1). For isolated objects such as the vault or ribosome, the peaks appear insular and sharp, while membrane or microtubules show elongated and continuous peaks consistent with the extended and repetitive character of the objects. Remarkably, TM identifies also low density and low abundance particles with high fidelity (Fig. 1). Automatic and semi-automatic particle detection algorithms have been widely tested for high-contrast and abundant macromolecular complexes in tomograms (e.g., ribosomes). However, fundamental macromolecular complexes such as the NPC or vault, which are scarce (2-3 copies per tomogram) and have low density, are particularly challenging. With optimal parameters, TM results in strong peaks for both macromolecular complexes (Fig. 1) and finds all positions identifiable by expert inspection. This finding is important in two ways: firstly, these complexes are fundamental for our understanding of cellular function, and secondly, given their low abundance, harnessing all the particles is key for the visual proteomics analysis.

### Optimization of TM parameters

Optimal TM requires systematic tuning of the bandpass filters (Fig. 2A,B), template (SI Figs. 1 and 2) and mask size (Fig. 2C), voxel size (SI Fig. 3) and angular sampling (SI Fig. 4). Optimal parameter values depend on the quality of the data as well as the size and shape of the object (Fig. 2D-F and SI Figs. 1 and 2). In tests on ribosomes, we found that mask tightness has a negligible effect as long as the template is completely contained (Fig. 2C). Different masking may also affect performance, e.g., with membranes included or excluded for membrane-associated structures (SI Fig. 5). Ribosome peaks decay sharply with increasing high-pass filter, i.e., when low resolution information is gradually removed (Fig. 2A). The low-pass filter has a less pronounced effect, although the z-score slightly increased when high resolution information was included (Fig. 2B). This analysis implies that, at least for ribosome detection, TM detection benefits from retaining high resolution information in the data.

**Fig. 2:**
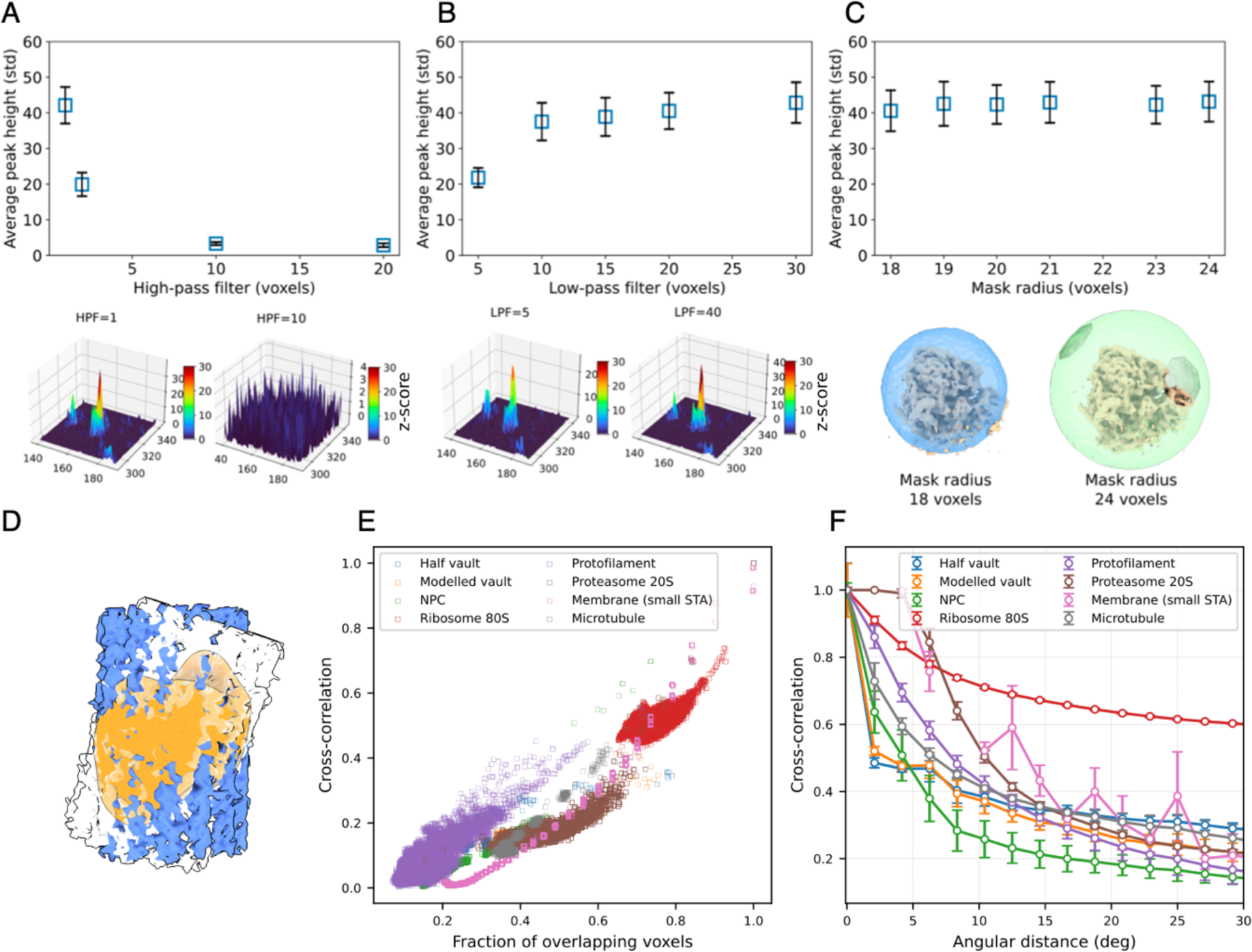
Optimization of the search parameters in template matching. A-C, Dependence of the average constrained cross-correlation peak height (z-scores) for 80S ribosomes on the high-pass (A) and low-pass filters (B), and on the diameter of the spherical mask (C). In A, no low-pass filter was applied, and in B no high-pass filter. The filters are shown in Fourier voxels (as defined in STOPGAP). D, Schematic representation of the overlapping voxels (orange) when a microtubule is rotated around its optimal orientation by 20 deg. E,F, Dependence of the constrained cross-correlation of a template with itself (*in silico* evaluation) as a function of the fraction of overlapping voxels) and the angular distance (F). In E, the CC values for all the rotations in a 10 degrees grid are shown. In all cases, error bars correspond to one standard deviation.

The impact of other parameters, such as voxel size or the number of orientations sampled, depends on the template mass, shape and size as well as the data. Therefore, we developed a Python-based tool that allows the user to perform an *in silico* evaluation of the TM parameters (see details in Methods and examples in SI Figs. 1 and 2). The *in silico* evaluation of multiple templates showed that the CC depends almost linearly on the fraction of overlapping voxels between the rotated template and the object (Fig. 2D,E). The number of overlapping voxels depends on both angular sampling and object shape (Fig. 2E). This effect is particularly pronounced for hollow objects such as the vault and elongated structures such as protofilaments. In such cases, even small rotations lead to a large decrease in the number of overlapping voxels and hence in the cross-correlation. Fig. 2E also shows objects that require finer orientation sampling to be localized with high confidence, requiring more computational power for detection. We conclude that general recommendations for sampling during template matching cannot be made. Therefore, our pipeline optimizes parameters *in silico* in a template specific manner, prior to analyzing experimental data.

### High-confidence TM accounts quantitatively for ribosome localizations

Reliable particle detection is a prerequisite for a quantitative analysis of the localization and interaction of molecular complexes. We assessed the ability of optimized TM to locate individual ribosome positions and orientations by comparing the results of TM with existing annotations of the cytosolic 80S ribosomes for *D. discoideum*^4^. The annotations were obtained in a multistep classification procedure using Relion^38^, as described in reference^4^, which resulted in a map with resolutions of up 4.5 Å.

Fig. 3 shows the results for TM on 4-binned data (8.704 Å/voxel). Motivated by our *in silico* evaluation (SI Fig. 4), we assessed the effect of the number of orientations by sampling the rotational space in angular steps of 30, 20, 10, and 5 degrees (576, 1944, 15192, and 119952 orientations) and selected TM peaks corresponding to local maxima in the z-score map that are above a threshold (Fig. 3). We considered a particle in the ground truth as “TM-detected” if it was located within 10 nm (∼1/3 of the ribosome diameter) of a TM peak. With increased numbers of orientations, the z-scores of the peaks increased and with that the percentage of TM-detected particles (Fig. 3C,D; see also SI Figs. 3 and 4). With orientations separated by ∼5 degrees, TM detected ∼95% of the 437 previously annotated particles with a mean distance to the TM peak of (3.73±1.57) nm and with orientations that closely matched the annotated orientations (Fig. 3F). Consequently, the averages of the particles detected and orientated by TM recapitulate the density of the 80S ribosome with high sensitivity and accuracy without the need for a multistep classification process (Fig. 3H, I). This suggests that TM can be used for a quantitative accounting of the particles present in the tomograms. Our analysis shows that the comprehensive search of the rotational space enhances the quantitative capability of TM in a trade-off with increased computational cost.

**Fig. 3:**
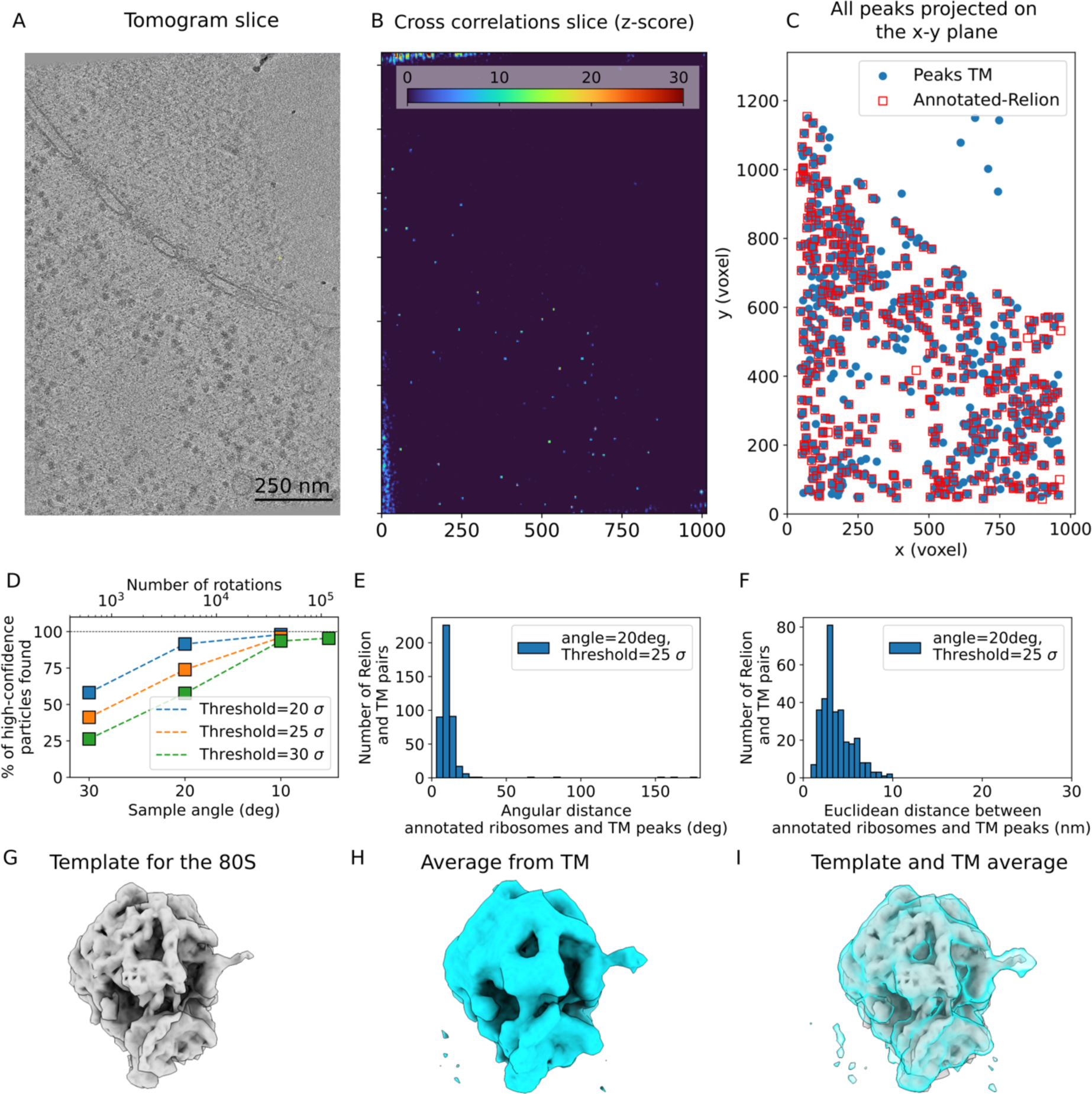
Template matching locates the 80S ribosome with high spatial and rotational accuracy. A, Tomogram slice (EMD-XXXX) showing abundant ribosomes. B, Slice of the z-score map obtained from template matching using the template of the 80S ribosome shown in G. C, Superimposition of the peaks obtained from template matching (blue circles; sampled every 5 degrees and with a cross-correlation threshold ≥30𝜎) and the high-confidence localizations obtained from an expert multiple-step alignment using Relion^38^ reported in reference^4^. D, Percentage of the high-confidence Relion particles detected within 10 nm of the TM peaks as a function of the rotational sampling (i.e., number of orientations). E,F, Histograms of angular (E) and Euclidean distances (F) from the TM peaks to the annotated Relion particles, respectively, each obtained for a 20-degree angular sampling. G, Template for 80S ribosome. H,I, Ribosome structure obtained by averaging the particles from TM (H, no further processing) superimposed on template (I).

### High-confidence TM reveals membrane compartments

Accurate segmentation of membranes is crucial for visualizing cellular landscapes. We tested TM for membrane segmentation with models of different origin and size (Table 1, Figs. 1 and 4). The first template was the map created from a frame in the trajectory of an atomistic simulation of a membrane in explicit water (atomic model). The second and third models were averages of the nuclear envelope obtained by subtomogram averaging with diameters of 43.5 nm (small STA) and 87 nm (large STA), respectively. For comparison, during TM, cylindrical masks with a diameter of 34.8 nm were used for both the atomistic and the small STA, while a cylindrical mask with a diameter of 76.5 nm was used for the large STA (see Methods).

The inner and outer membranes of the nuclear envelope were detected using any of the three templates (atomistic, small STA, large STA; see supporting video). The atomistic and small STA templates performed roughly on par. Increasing the number of orientations (20, 10, and 2 degrees at 4-binned data with 8.704 Å/voxel) consistently decreased the background noise (Fig. 4), sharpening the peaks, and increased the confidence in the TM detection. False positives for the small templates (atomistic, small STA), e.g., from a microtubule segment (Fig. 4 left; see also Fig. 1) are suppressed by using the large STA template (or, visually, by recognizing the lacking 2D extension). However, the large STA model gives only a weak signal for curved membranes, pointing to the need for an expanded model set of membrane patches of varying curvature.

**Fig. 4:**
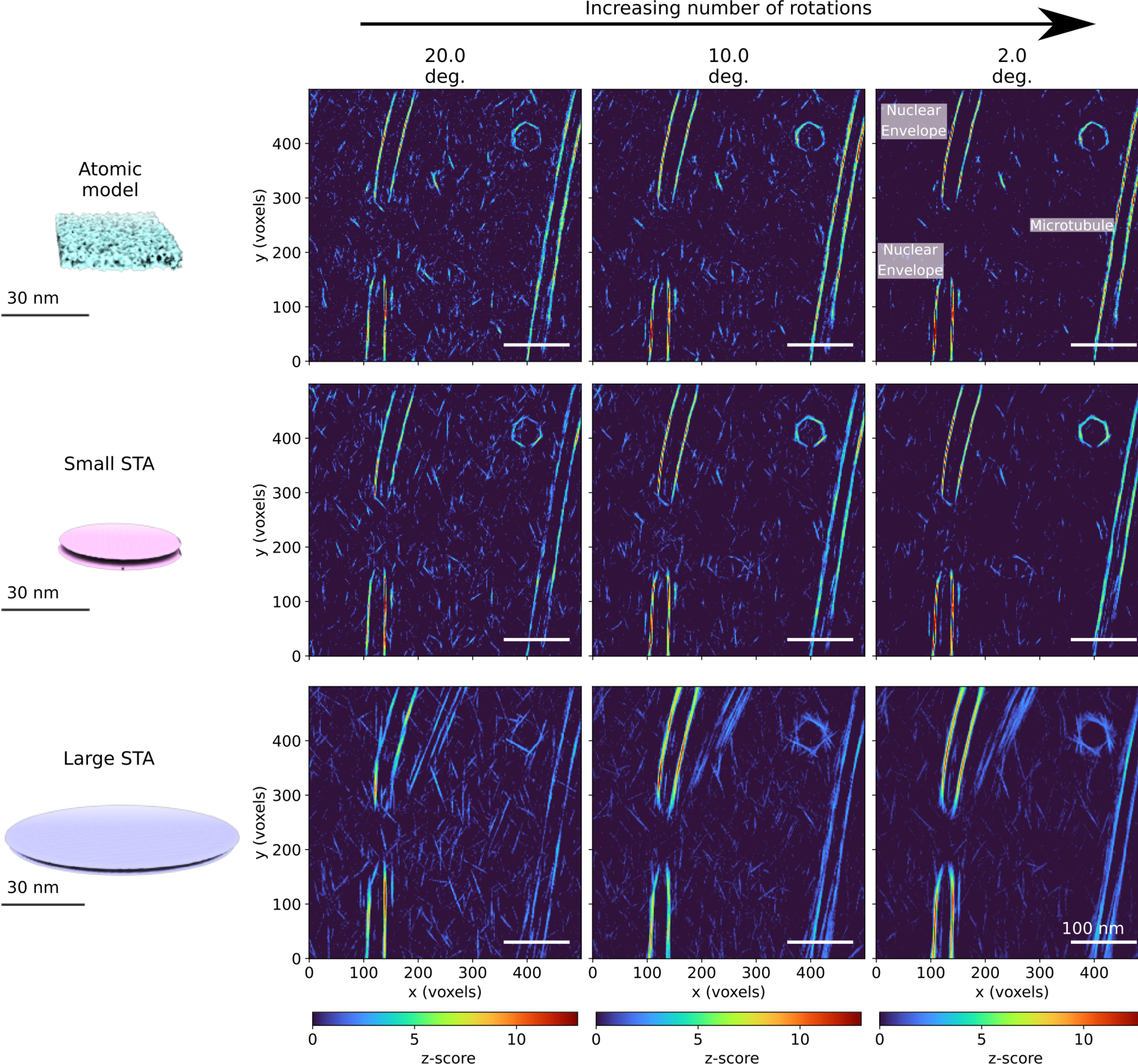
Template matching for the segmentation of membranes in 3D. Results in the top, middle and bottom row were obtained for templates constructed from a simulated atomic membrane and STAs of the nuclear membrane with diameters of 50 (small STA) and 100 (large STA) voxels, respectively (8.704 Å/voxel). The results from left to right correspond to increasing angular sampling of 20, 10 and 2 deg, respectively. Note that the peaks at the upper right corner originate from a highly curved vesicle while the two stripes on the right-hand side of the cross-correlation maps (z-scores) are a microtubule and not a membrane (see Fig. 1).

Although computationally expensive compared to other segmentation methods, template matching for membranes has several strengths. For example, the template matching output could be used as an initial annotation for training deep learning algorithms. In addition, TM not only predicts the positions of the membranes in the tomogram, but also provides voxel-by-voxel normal vectors, which in turn enables a detailed analysis of the local properties of the membranes. The latter could also be used as an automatic input for triangulation methods and/or as a starting point for simulations of membrane dynamics.

### TM locates individual subunits down to 100 kDa mass

We tested the ability of TM to localize subunits and assign substates of ribosomes, the NPC, and microtubule fragments. We generated templates for the subunits of the *D. discoideum* NPC according to its C8 symmetry, microtubule protofilaments, the small (40S) and large (60S) ribosomal subunits, and for two prominent 80S ribosome states capturing the ratchet-like motion essential for protein synthesis^39^.

For the ribosomal subunits, we performed TM on 2-binned data (4.352 Å/voxel) with orientations every 10 degrees, since TM on 4-binned tomograms showed inconclusive peaks. A sub-volume of the tomogram was analyzed independently with three different templates: 80S, 60S, and 40S (Fig. 5). Similar to the 4-binned data (Fig. 3), the TM localized 96.9% of the 80S annotated ribosomes with CC peaks up to 114 𝜎 (Fig. 5B,C). Furthermore, when comparing the positions and orientations of the subunits, TM correctly predicted the location of the subunits and their relative orientations (Fig. 5D). Small but noticeable differences between the orientations of the subunits with respect to the position of the 80S reflect the limited angular sampling.

**Fig. 5:**
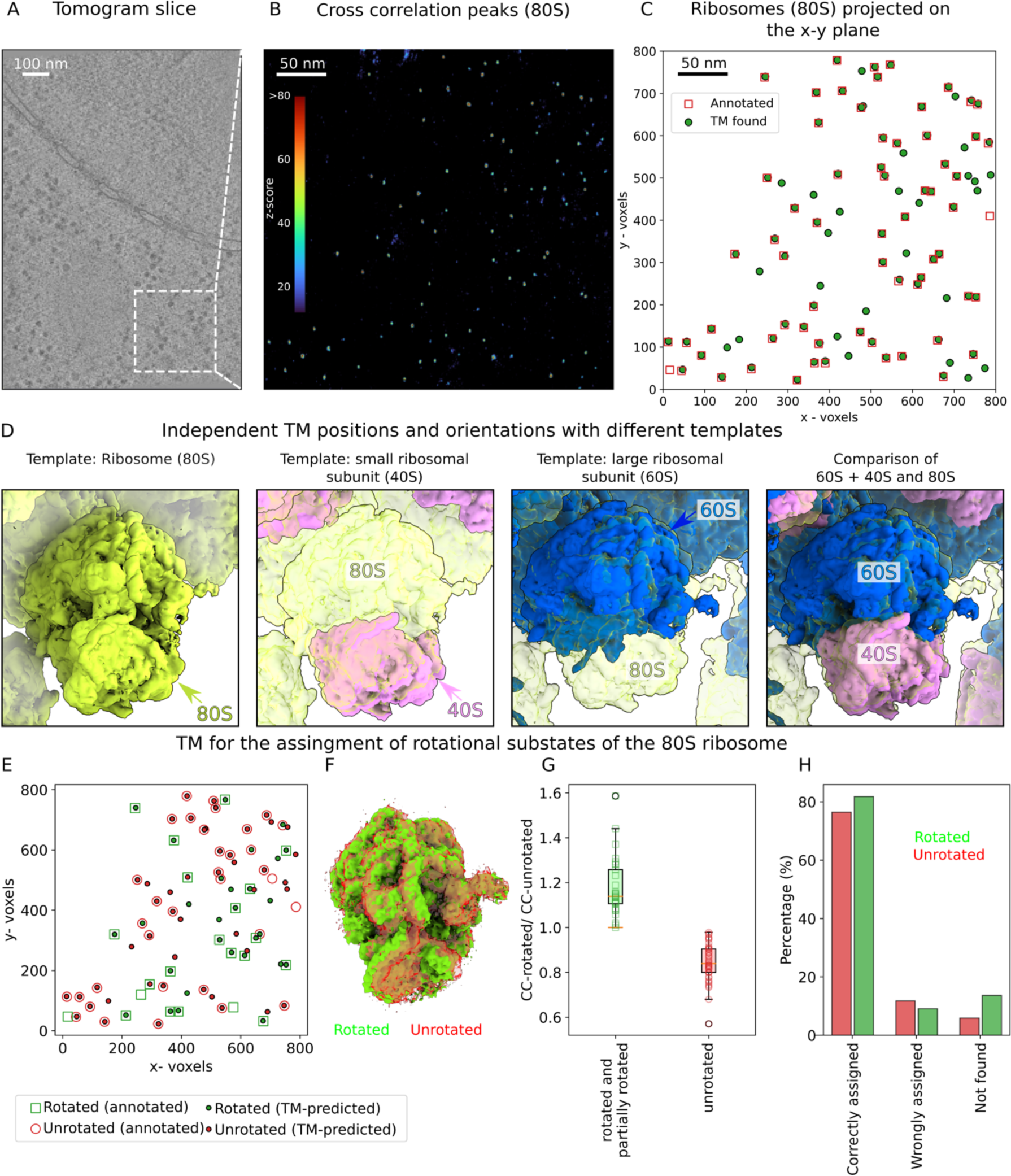
Template matching predicts the relative orientations of ribosome subunits and assigns ribosome rotational States. A, Tomogram slice. B,C, TM finds the positions of the 80S (green) ribosomes annotated with high confidence (96.9% of particles within 2.4±1.5 nm). Two more templates were tested, (i) the small (40S - pink) and (ii) the large (60S - blue) ribosomal subunits. D, Predicted position and rotation of the 40S and 60S compared to the predicted orientation for 80S from TM. Note that the positions and orientations displayed the subunits and the 80S were obtained from independent TM calculations. E, Comparison of the assignment of the ratchet-like rotational states of the ribosome from TM using a Gaussian mixture model^40^ with existing annotations^4^ as function of position. F, Rotated and unrotated template. G, Ratio of TM scores as function as function of assigned rotation state (line: median; box: interquartile range; error bars: Range). H, Consistency of assignment (annotated as rotated: red; unrotated: green).

Directly from the z-score maps, we could deduce the canonical 8-fold symmetry of the NPC (Fig. 6A) after performing TM on 4-binned tomograms (8.704 Å/voxel) and sampling orientations every 10 degrees as suggested by our *in silico* analysis. Two NPCs at the edge of the lamella have only 7 (left) and 5 (center) detectable subunits left after the milling process. Interestingly, no peaks were detected using an NPC from a different species as a template (SI Fig. 5).

**Fig. 6:**
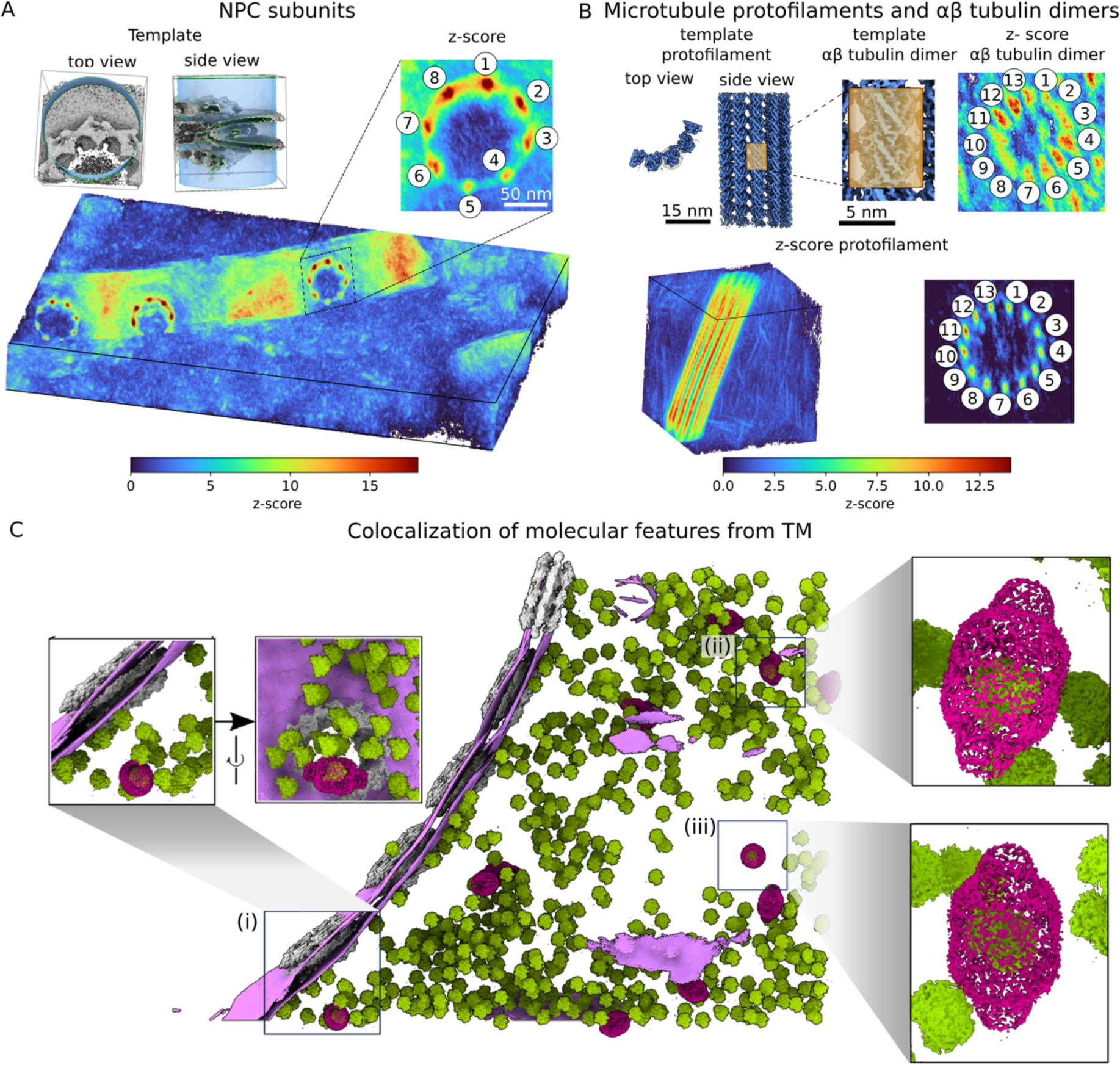
Template matching detects NPC subunits, microtubule protofilaments and ribosome-loaded Vaults. A,B, Perspective view of the three-dimensional constrained cross-correlation map obtained from template matching using an NPC subunit (A) and microtubule section (B). The templates are shown in the upper left-hand corner of each panel. The 8- and 13-fold symmetries for the NPC and the microtubule, respectively, emerge naturally from template matching (zoom-ins with numbered peaks). The NPC subunit and protofilament templates were cut from the whole NPC and microtubule templates, respectively. The αβ-tubulin dimers in B were obtained by masking the protofilament template (orange box). C, From the three-dimensional localization maps generated for visualization^58^ and analysis, TM finds ribosomes inside vaults indicated by the squares (i, ii and iii clockwise from left). In (i), the vault (magenta; 54𝜎) containing the ribosome (green; 63𝜎) is near a NPC (gray) and the nuclear envelope (purple). In (ii) and (iii), the vaults (32𝜎 and 59𝜎) containing the ribosomes (46𝜎 and 77𝜎) are in the cytoplasm.

In a high-resolution map (2-binned, 2.446 Å/voxel) and with fine angular sampling (5 degrees), TM resolved individual αβ-tubulin dimers as distinct peaks with the 13-fold symmetry of microtubules (Fig. 6B). Despite the low combined mass of only 100 kDa, TM achieves good statistics both in terms of true positives and (likely) false negatives, using a protofilament fragment as reference (SI Fig. 6). SI Fig. 7 further examines the gain in matching confidence for microtubule fragments at lower resolution (8.704 Å/voxel, 10 degrees).

### TM identifies functional substates

By comparing the relative TM z-scores on 2-binned data (4.352 Å/voxel) with orientations every 10 degrees, we could correctly assign the ratcheting state of the small subunit of individual ribosomes in space (Fig. 5E-H). Two known representative ratcheting states of the *D. discoideum* ribosome were used as templates^4^: rotated (emd-15815) and unrotated (emd-15812), and the states were assigned using the expectation-maximization algorithm (see Methods for details) to predict the mixture of subpopulations (Fig. 5G), similar to previous studies^40^. Although the rotated and unrotated templates share most of the density with only a slight rotation of the 40S (Fig. 5F), the TM assignments differentiated between the rotated and unrotated states, matching the existing annotations in 77.7% and 82.4% of cases, respectively (Fig. 5E,H). It is worth noting that there are other intermediate rotation states, and the binding of multiple cofactors to the ribosome along the translation cycle^4^ may affect the TM z-scores and ultimately the state assignment, which may account for non-matching particles.

Overall, these results demonstrate that TM can find subunits of macromolecular complexes with high accuracy and precision. The peaks we obtained for NPC subunits, microtubule protofilaments, and ribosomal subunits with our TM approach recover known features of the particles previously identified by a variety of structural and biochemical approaches^4,39,41,42^. For the 80S ribosome, the TM analysis spatially assigned known functional states of translation with good accuracy. The ability to identify individual subunits or conformational states within a given tomogram maximizes the use of available information for structural purposes and can lead to the discovery of non-canonical symmetries. By chance, one may in this way also capture complex assembly intermediates.

### High-confidence TM identifies vault encapsulated ribosomes in situ

The biological function of the vault particle remains mysterious. A few interactors binding to the inside surface have been reported^43,44^ which in line with its capsule-like morphology has led to speculations that vaults may enclose other particles and transport cargo within the cell. To the best of our knowledge, however, evidence for vaults encapsulating cargo *in situ* is yet missing. Three of the vaults in the tomogram of Figs. 5A and 6C contain 80S ribosomes with highly significant z-scores (vaults: 54𝜎, 32𝜎, and 59𝜎 ribosomes: 63𝜎, 46𝜎, and 77𝜎, in Fig. 6C(i)-(iii), respectively). These findings support the hypothesis that vaults can be cargo-loaded *in situ*. Whether the encapsulation occurred during vault biogenesis or by transient opening remains to be further investigated.

## Discussion

The comprehensive identification of particles in electron tomograms remains challenging. Despite its conceptual simplicity, template matching has been considered a low-precision method, and its application has been limited by the low signal-to-noise ratio of tomographic data, the scarce availability of suitable templates, and the lack of objective optimization of search parameters. Here, we have shown that template matching can identify the positions and orientations of multiple macromolecular complexes in living cells with high accuracy and fidelity. For this task, templates can be used from multiple sources such as data banks, simulations, homology modeling, or volumetric data from the tomograms. For maximum efficiency, we developed an *in silico* parameter optimization software.

With optimized TM, we achieved a mass resolution of 100 kDa in experimental tomograms of a crowded cell. Using a generic template for human tubulin, we could readily localize individual αβ-tubulin dimers in a high-resolution CryoET map of *D. discoideum* cells (Fig. 6B, SI Fig. 6). High-confidence TM thus pushes into a particle size regime *in situ* that covers much of cellular biology.

By exploiting geometric and contextual features, one can further improve the likelihood of finding objects by template matching. Vaults, for example, are low-density objects, but their unique shape facilitates identification with confidence (Fig. 1 and SI Fig. 1). Spatial extent is also important for TM. Another strategy to search for smaller objects is to decrease the voxel size (increase magnification), which in this case allowed us to locate ribosomal subunits or distinguish between ribosomal substates. However, the location of smaller isolated objects poses an additional challenge: the unambiguous validation of the peaks. In the cases presented here, we used annotated data (Figs. 3, 5 and SI Fig. 3) and expert inspection (Figs. 1, 4, 6 and SI Figs. 5, 7 and 8). However, when considering smaller structures, an increase in the number of peaks and volumes of high significance is expected. To overcome this challenge, we envision a hierarchical approach in which we mask parts of the volume where we have a high confidence of the presence of an object, and then perform a focused search for smaller objects. Still, relying on template matching alone may be insufficient and additional information, such as abundance data would need to be incorporated to effectively analyze TM results.

The widespread use of template matching is still limited by its computational expense, due to the nature of the algorithm that evaluates each voxel in the volume for up to hundreds of thousands of orientations. This problem is exacerbated in the current STOPGAP implementation with limited parallelization across CPUs. We see significant potential for improvement by exploiting the parallel capabilities of graphics processing units (GPUs) as the TM algorithm is by construction embarrassingly parallel and relies on Fourier transformation, which is highly efficient on GPUs.

Finally, the TM workflow can readily be combined with AI-based approaches^22,28–31^. At one end of the pipeline, AI can be used to optimize TM parameters and, at the other end, to integrate the outputs across template families into classification scores. At the center of the pipeline, however, the 3D CC score is highly efficient and captures the relevant physics by being rigorously proportional to the log-likelihood for Gaussian noise in the 3D map^45^ (Methods). In future, TM-annotated tomograms can be used to train and validate AI-based particle localization methods.

Taken together, our analysis demonstrates the detection of various objects, with high confidence, in cryo electron tomograms acquired with the latest hardware. By expanding the repertoire of templates, e.g., from AlphaFold^46^ and molecular dynamics simulations, TM should help us assign molecular identities to the large parts of tomograms currently unassigned. High-confidence TM thus changes the workflow in CryoET through fast, automated, objective and comprehensive feature identification. In turn, CryoET combined with high-confidence TM brings us closer to the goal of visual proteomics: to map the positions and orientations of all macromolecular complexes within living cells.

## Methods

### Experimental tomograms

All tilt series of *D. discoideum* used in this study, as well as the annotations for the ribosomes and their substates were previously reported^4^. The cell culture, sample preparation, data acquisition, and image processing are detailed in the original publication. In brief, tilt series were collected at 300 kV on a Titan Krios G2 microscope equipped with a Gatan BioQuantum-K3 imaging filter in counting mode and a Titan Krios G4 microscope equipped with a cold FEG, Selectris X imaging filter, and Falcon 4 direct electron detector in counting mode. Projections had a pixel size of 2.176 Å and 1.223 Å, respectively, and were acquired in a dose symmetric acquisition scheme^47^ with 2 deg increments^4,48^. The initial tomogram reconstruction was performed in eTomo from IMOD^49^ and the established parameters were used to reconstruct the tomograms with 3D-CTF correction using novaCTF^50^. The corrected tomograms were used for TM either in their unbinned form or with applied binning of 2, 4 or 8.

### Template matching produces maximum-likelihood solution

For Gaussian noise of width 𝜎 in the intensities of a CryoET 3D map 𝑀, the likelihood 𝐿 that a feature in the map is consistent with a template 𝑇 rotated and translated by 𝑅 is proportional to

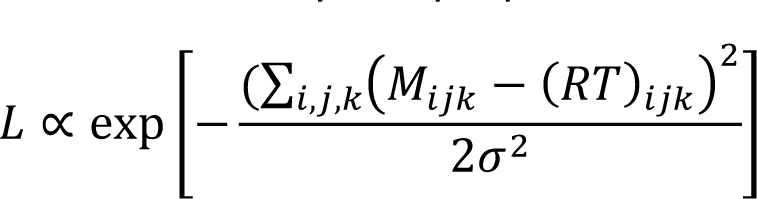

By multiplying out the square, summing over the voxels 𝑖, 𝑗, 𝑘, and recognizing that the “𝑀^$^” and “(𝑅𝑇)^$^” terms are constant, we find that

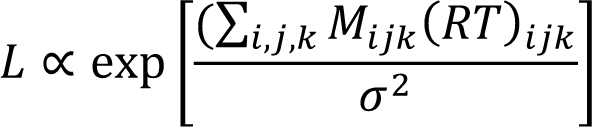

The term in the exponent is exactly the cross-correlation CC between map and template divided by 𝜎^$^. Analogous to single-particle 2D images^45^, the cross-correlation of template and map is thus the log-likelihood scaled by the squared noise amplitude. CC optimization over template rotation and translation 𝑅 thus gives the maximum-likelihood solution.

### Template matching

We performed TM using STOPGAP^35^ for the cases described in Table 1. STOPGAP is an open-source freely available Matlab-based code: https://github.com/williamnwan/STOPGAP. As input, STOPGAP requires a template, a list of orientations to probe (angular sampling), a “wedge list”, definitions of the filters, and the reconstructed tomogram. Details on the preparation of the templates are given below. The list of orientations was generated using STOPGAP function generate_angle_list, which samples the angle space uniformly on a grid and also takes into account the symmetry of the template (see Table 1). The wedge list contains the acquisition parameter information for the individual tilts. In particular it must contain: the pixel size, the tilt angle, defocus and electron dose. The low-pass filter allows low-frequency signals to pass through while attenuating high-frequency signals, while high-pass filters allow high-frequency signals to pass through while attenuating low-frequency. In STOPGAP, both are defined in voxels defining the radius of a spherical mask applied in Fourier space (i.e., these values depend on the dimensions of the template). See further details in the STOPGAP documentation. For each template, we obtained a map of the maxima CC of the local cross-correlation over orientations, which we turned into z-scores as 𝑧 = (CC − 𝜇)/𝜎 with 𝜇 and 𝜎 the average and standard deviation of CC values across the map, respectively.

### *In silico* peak analysis

The template weighting and CC calculation methods from STOPGAP^35^ have been ported to Python and extended to output additional information relevant to the input parameters. The inputs are the same as for the original TM, but instead of a whole tomogram, another small volume is used. The volume can be either the same as the template (typically an STA map or a model) or a subtomogram (obtained either based on an existing ground truth or by manual picking). For full peak analysis, one must also provide a density mask, a binary map corresponding to the density of the template (or alternatively a threshold to create one during the analysis). In addition to the z-score map and the angles map, the peak analysis provides additional information from the TM progress as well as the analysis of the template and the resulting maps. From the TM progress, it outputs a table showing the dependence of the template orientation on the CC scores and on the number of overlapping voxels. For the template, it computes the dimensions, the number of voxels in the density mask, and a solidity calculated as the number of voxels in the density map divided by the volume of its convex hull. It also returns the value of the peak, its exact location, and line profiles through the peak along each dimension. The angle map is used to compute three maps of angular distances, where each voxel contains the angular distance in degrees between the orientation encoded in the angle map and the starting template orientation. The first map contains the angular distance of the full orientation and is computed using a quaternion-based cosine similarity formula^51^. The second map contains the angle between the normal vectors of the final and the initial orientation, which encodes the rotation on the cone. Complementarily, the third map contains the angle between the in-plane vectors. The maps provide information on whether the CC scores are more sensitive to cone or in-plane rotation (or neither), and thus can be used to determine sufficient angular sampling. Finally, the key results of the peak analysis are summarized in a PDF file to provide an easy-to-read overview for the user. While the tool is most useful for determining the optimal setup for STOPGAP TM (or deciding its feasibility), it can also be used to analyze the origin of false positive results by testing a template against a map containing a different structure. For example, a ribosome template can be tested against map containing proteasome to determine the pixels size and filtering to distinguish these two with sufficient confidence. Similarly, the membrane template can be used against a microtubule structure to determine size of the template and mask necessary to pick mostly membranes. Lastly, there is a possibility to turn off the missing wedge weighting to analyze its impact on the peak shape or add an angular offset to the starting orientation to see how it affects the peak value for given angular sampling.

### Membrane templates

For the “atomistic” membrane template, we used the final lipid bilayer of a 28-ns molecular dynamics simulation of a 40×40 nm^2^ membrane patch in explicit water, using the setup and protocol of reference ^52^. The “small STA” and “large STA” models were obtained as subtomogram averages of the nuclear envelope with diameters of 43.5 nm and 87 nm, respectively. In TM, cylindrical masks with a diameter of 34.8 nm were used for the atomistic and small STA models. For the large STA model, the diameter was increased to 76.5 nm.

### Creation of density maps from atomic models

In the cases where an atomic model was available (membrane and vault), we used the molmap function of ChimeraX ^53^ with a resolution of 3.5 Å. Here, each atom is represented by a three-dimensional Gaussian. The width of the Gaussian is given by the resolution, while the amplitude is proportional to the atomic number. Afterward, we used EMAN2^54^ to rescale and resample the map and have identical voxel size as in the tomograms.

### Templates for the ribosomal subunits 40S and 60S

To generate templates of the large 60S and small 40S ribosomal subunits, the ribosome structure of a translating *D. discoideum* ribosome from EMD-15810^4^ was segmented in ChimeraX^53^ using the Segger function^55^. A fitted eukaryotic ribosome atomic model (PDB-id: 5LZS^56^) was used to guide this procedure.

### Statistical assignment of ribosomal substates

To assign the sub-states of the ribosome 80S, we performed TM using two templates corresponding to a rotated (emd-15815) and unrotated ribosomal states (emd-15812). High-confidence peaks were extracted from each TM map with their respective z-scores.

For a given particle, defined by its coordinates, we computed the ratio between the two TM z-scores as:

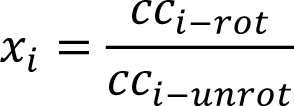

where 𝑐𝑐_*i–rot*_, 𝑐𝑐_*i–unrot*_ corresponds to the z-score obtained for the particle *I* with the rotated and unrotated template, respectively. To assign the rotational substates of each particle, we used gaussian mixture model (GMM). Specifically, we assumed that the distribution of ***x*** can be modeled as a linear superposition of two gaussian distributions, one for the rotated state and the other for unrotated state. The probability density function of the GMM can be written as:

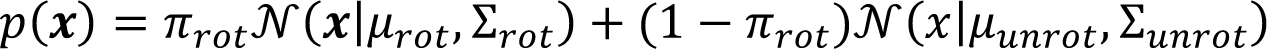

where 𝒩(𝜇, Σ) represents a gaussian probability density function with mean 𝜇 and variance Σ.

To estimate optimal parameters for 𝜋_*rot*_, 𝜇_*i*_, Σ_*i*_, we used the *sklearn.mixture*^57^ python implementation of the estimation-maximization (EM) algorithm. The EM algorithm alternates between computing the expected values of the latent variables (the assignment of each data point to a mixture component) and updating the parameters of the GMM to maximize the log-likelihood of the observed data. Specifically, the E-step computes the posterior probability of each mixture component for each data point, given the current estimates of the parameters, while the M-step updates the parameters to maximize the expected complete log-likelihood of the data, given the posterior probabilities.

## Data availability

The previously published structures for the NPC subunits (*Homo sapiens*) EMD-14325, EMD-14328 and EMD-14330, the 80S ribosome (*D. discoideum*) EMD-15810, EMD-15812, and EMD-15815, the 20S proteasome (*Homo sapiens*) EMD-4877 and the microtubule (*Homo sapiens*) EMD-6351 are accessible through the Electron Microscopy Data Bank. The previously published structures 7R5J, 6RGQ, and 3JAR are available through the Protein Data Base. The remaining EM densities and tilt series used in this study will be deposited the Electron Microscopy Data Bank upon publication.

## Code availability

All the code used for this study is part of the public repository: https://github.com/turonova/cryoCAT

## Acknowledgments

This work was funded by the Max Planck Society and the Chan Zuckerberg Initiative for Visual Proteomics Imaging. The Max Planck Computing and Data Facility is acknowledged for computational resources. We thank Stefanie Böhm for critical reading of the manuscript and Sonja Welsch and Iskander Khusainov for fruitful discussions. We also thank Martin Simonovsky for helpful discussions on peak analysis, Jürgen Köfinger and Jakob Bullerjahn for discussions on the Gaussian mixture model and Agnieszka Obarska-Kosinska for the help with the template of the human NPC subunit.

## Supplementary Information

**SI Fig. 1:**
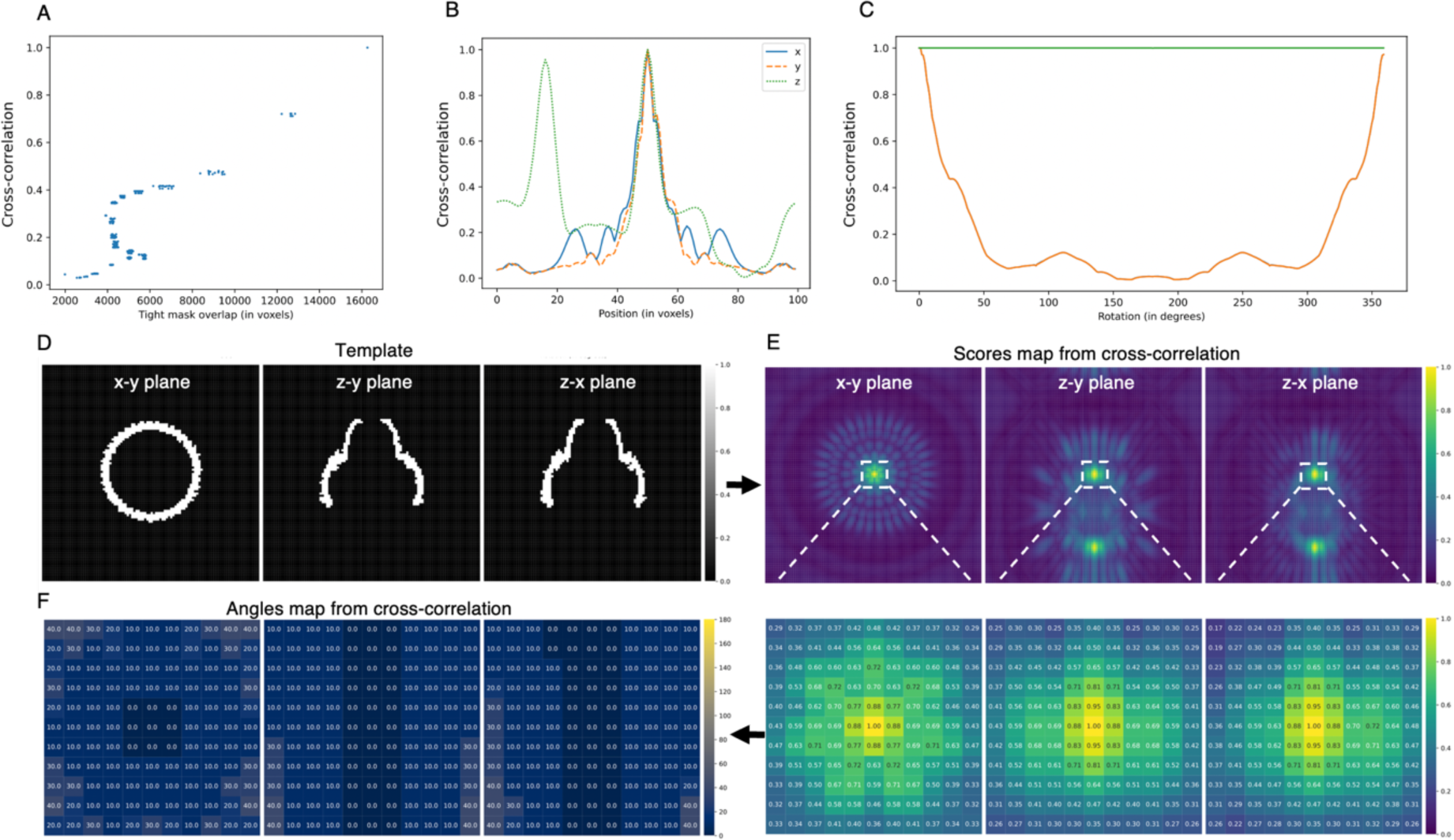
*In silico* assessment of cross-correlation for the vault. A-C, Cross-correlation as a function of the number of overlapping voxels (A), the position (B), and the angular distance (C). D, Template. For the vault, we used its two halves as template, resulting in two peaks along the z-direction. E, Scores-map (as in STOPGAP) with a zoom-in onto the peaks (bottom). F, Angles map from zoom-in onto cross-correlation.

**SI Fig. 2:**
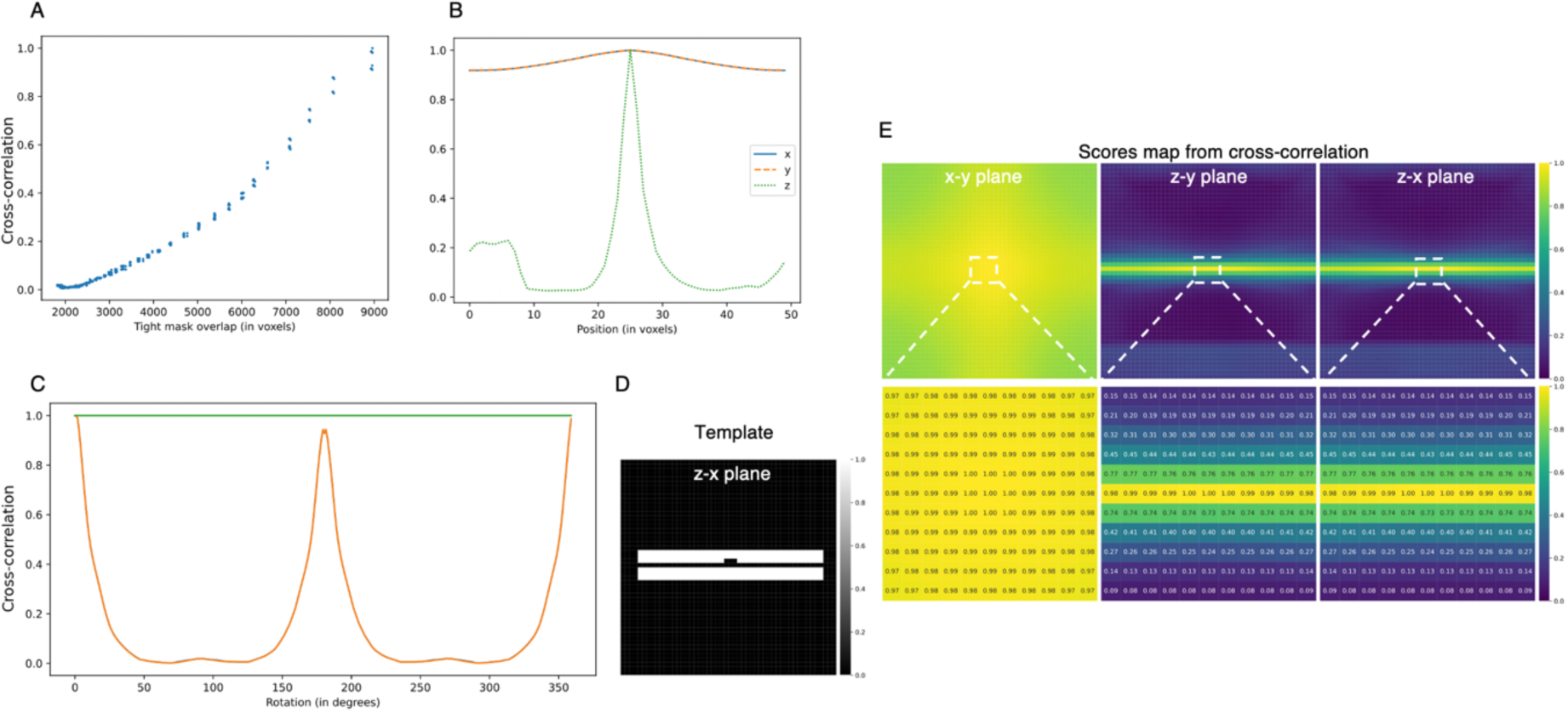
*In silico* evaluation of the cross-correlation for the large STA membrane. A-D, Dependence of the cross-correlation on the number of overlapping voxels (A), position (B), angular distance (C) and the used template (D). E, The CC scores map revealed a peak that extends into the x-y plane and was unaffected by rotation in the x-y plane, consistent with the symmetry of the membrane.

**SI Fig. 3:**
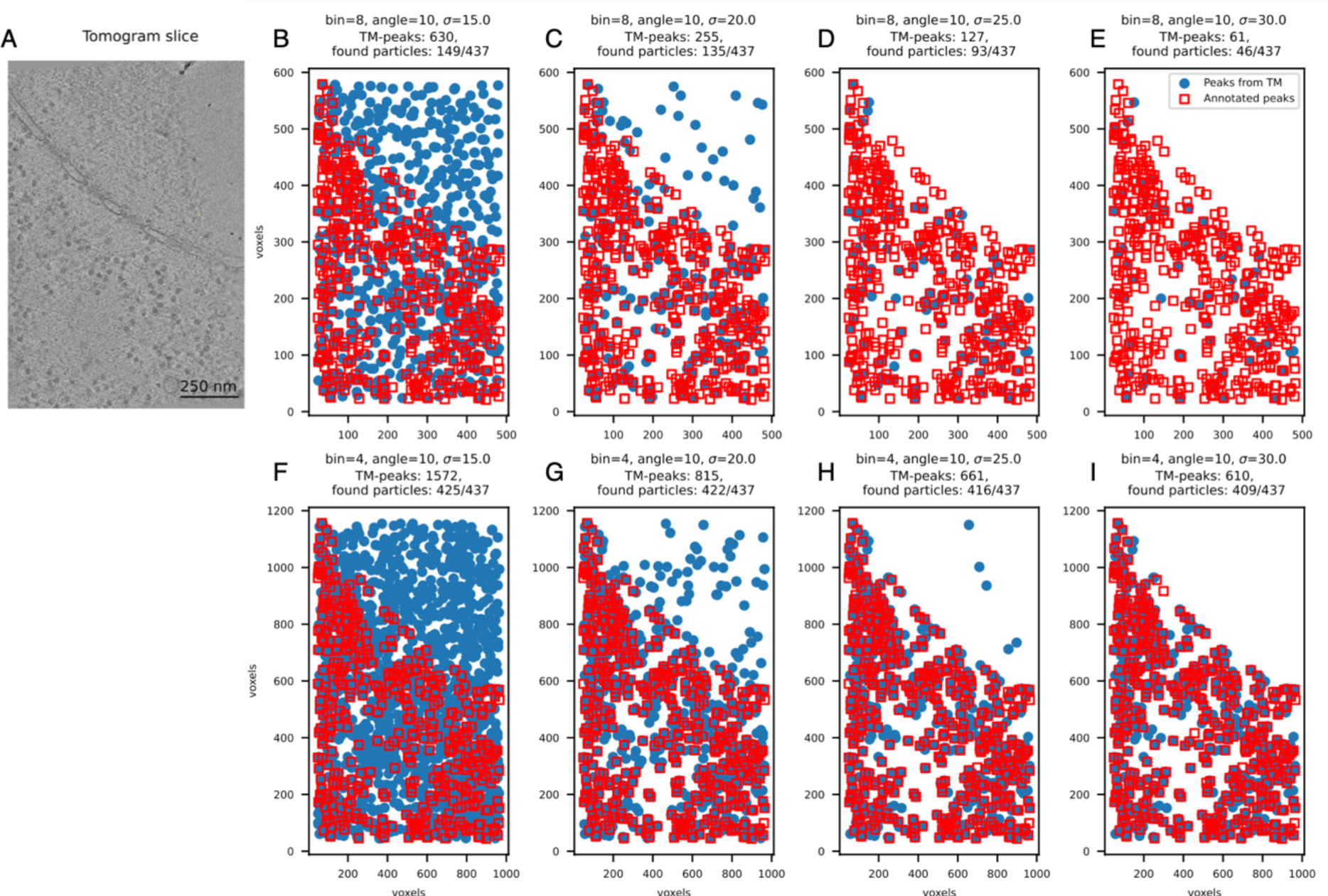
Template matching results for the 80Sribosome with 8-binned data (bin-8: 17.40 Å/voxel, top row) and 4-binned data (bin-4 of 8.704 Å/voxel, bottom row). A, Tomogram slice. B-I, Superimposition of the peaks obtained from template matching (blue circles; sampled every 10 degrees with varying cross-correlation thresholds) and the high-confidence localizations obtained from an expert multiple-step alignment using Relion^38^ as described in reference ^4^. The number of peaks as well as the ratio of “TM-found” particles is described in the figure.

**SI Fig. 4:**
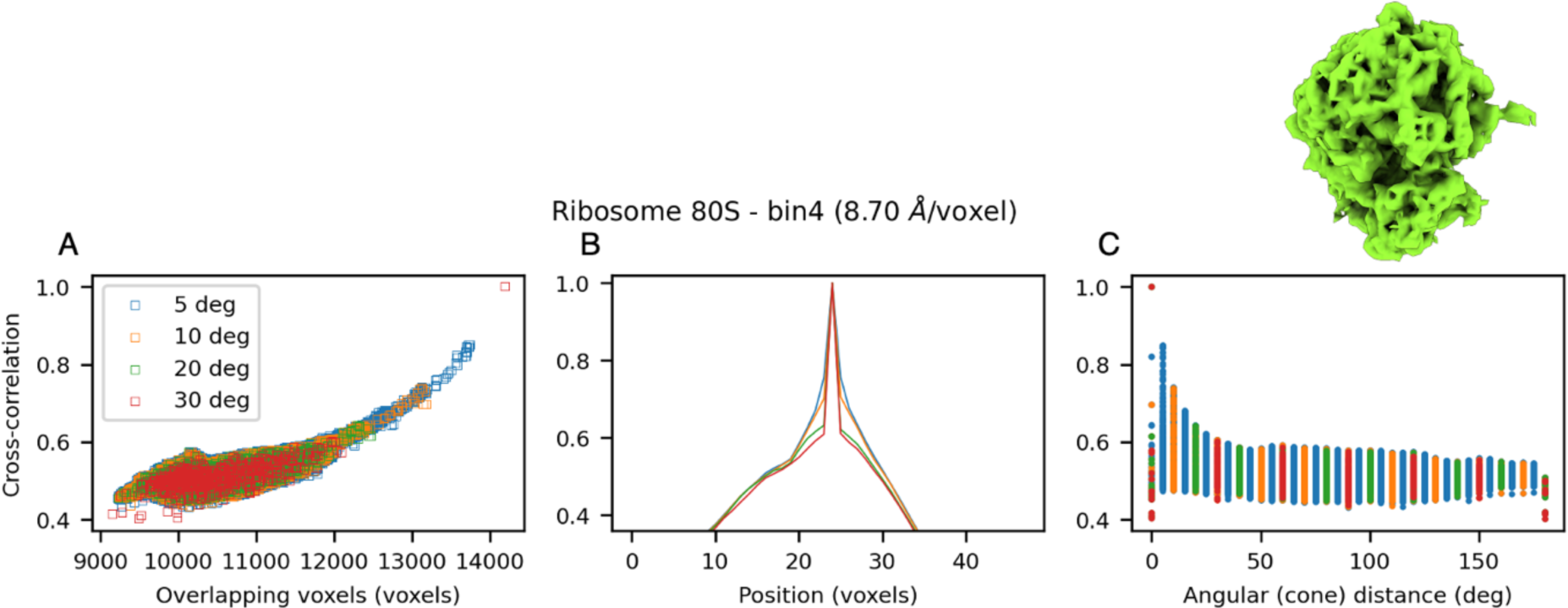
Evaluation of increasing the number of orientations in template matching of ribosome 80S using the python tool for *in silico* evaluation. A, Cross-correlation as a function of the absolute number of overlapping voxels for all evaluated rotations. B, Cross-correlation as a function of the distance along the z-plane. C, Cross-correlation for all evaluated rotations as a function of the angular cone distance. With an increasing number of orientations (decreasing angular sampling), more rotations lead to a higher number of overlapping voxels and thus to a higher cross-correlation. This effect leads to a broadening of the peak in the middle panel.

**SI Fig. 5:**
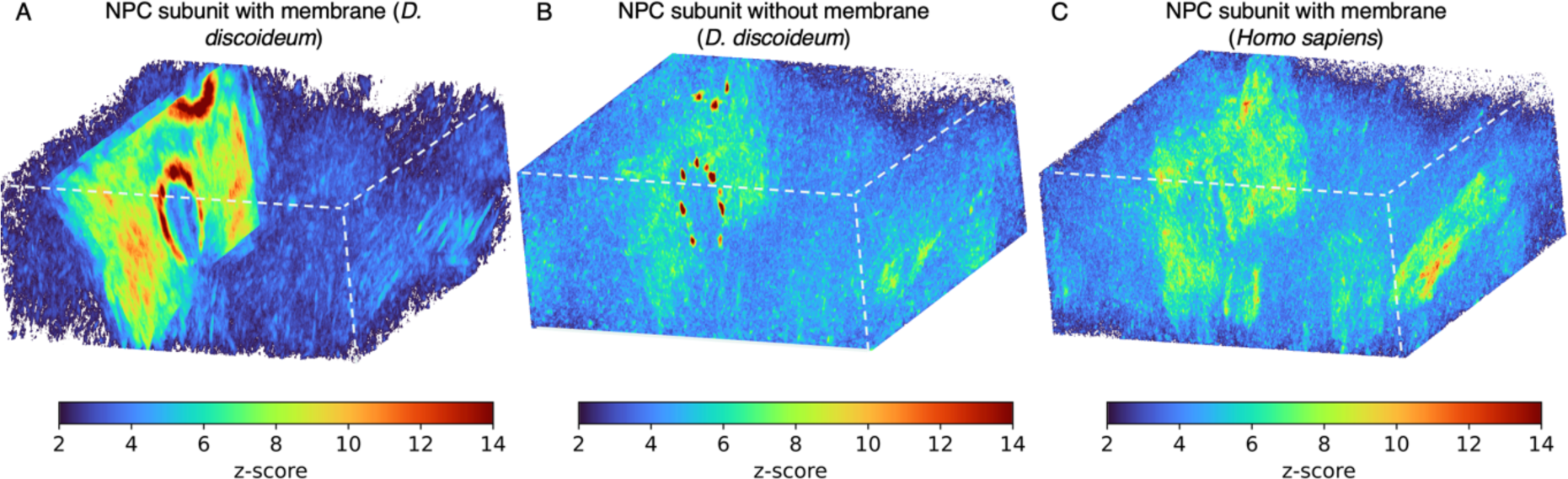
Template matching results for the NPC subunit of two different species. Cross correlation (z-score) maps obtained on the same tomographic volume using different templates of the NPC subunit. A-C, z-scores for NPC subunit templates with (A) and without membrane (B) for *D. discoideum*, and (C) with the NPC template for *Homo sapiens* NPC with membrane^12^. Note that no clear peaks are detected for the *Homo sapiens* NPC subunit, which exemplify the potential of TM for comparing macromolecular complexes of different species.

**SI Fig. 6:**
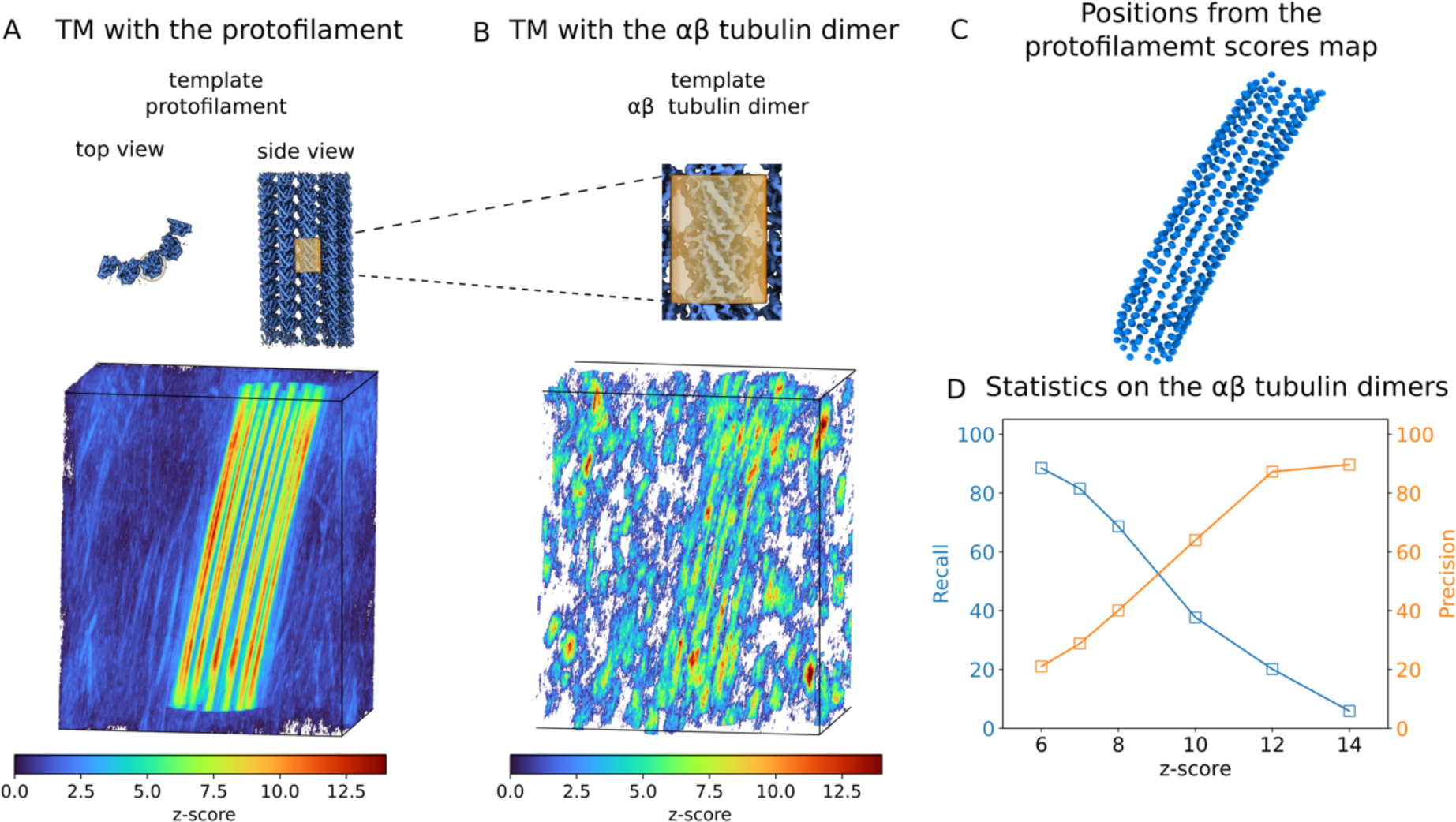
Statistics for template matching using a single ∼100 kDa αβ-tubulin dimer. A,B, Cross correlation (z-score) maps obtained on the same tomographic volume using a template containing four microtubule protofilaments (A) and a single αβ-tubulin dimer (B). c, Positions of the individual subunits extracted from (A), and adopted as annotated particles, correspond to local maxima in the z-score map, above 6𝜎, with a minimum separation of 14 voxels. D, Statistical analysis of the αβ-tubulin dimer localizations in the volume shown in B with the positions in C as reference. Shown are the recall = #(true positives) / [#(true positives) + #(false negatives)] (blue, left axis) and the precision = #(true positives) / [#(true positives) + #(false positives)]) (orange, right axis) as functions of the threshold applied to the tubulin dimer z-score. A position was considered as true positive if it was within 3 nm of a position in (C).

**SI Fig. 7:**
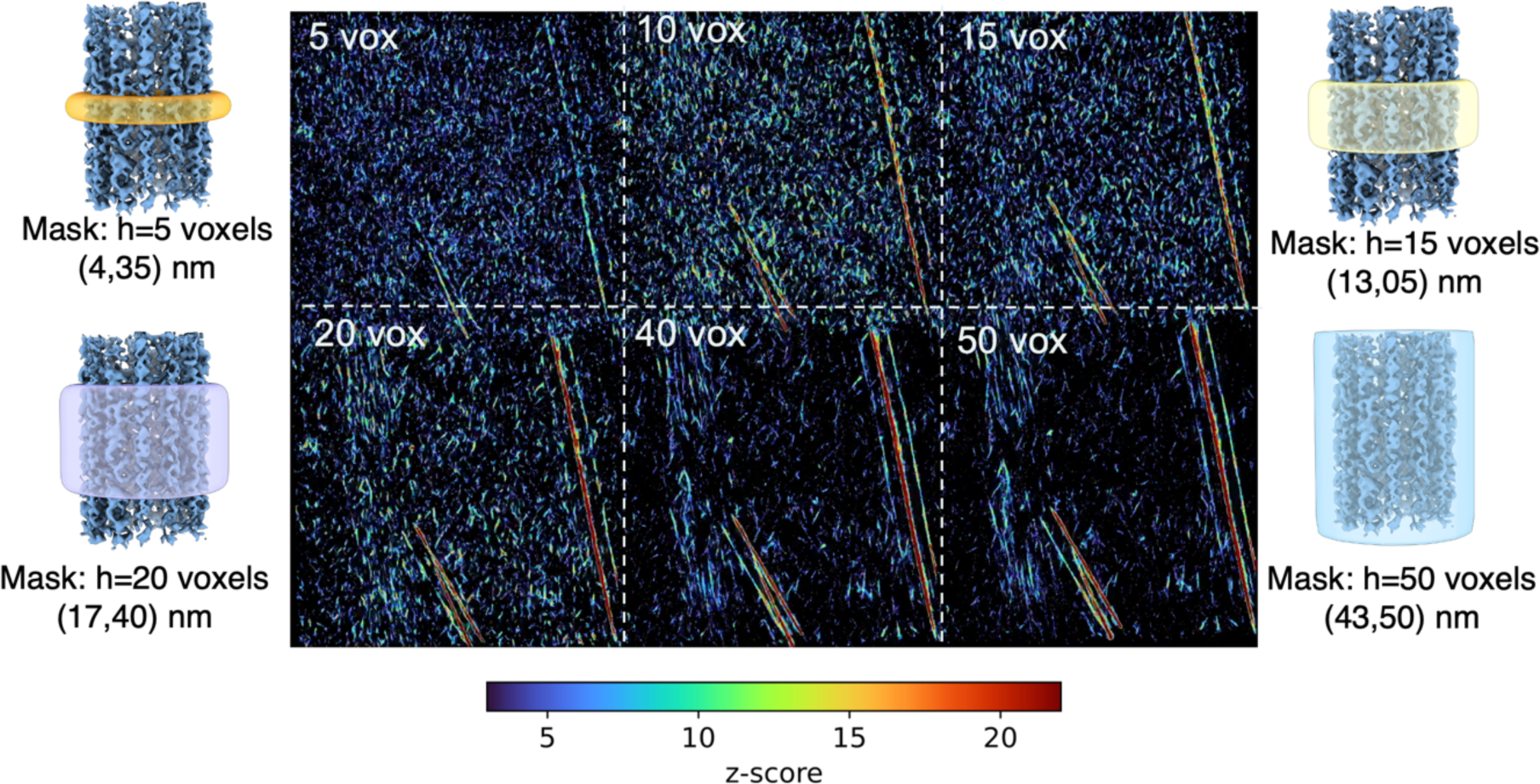
Effect of the effective template size on matching. The left and right panels show the microtubule template (center) with masks of different heights from 5 to 50 voxels (transparent outline). Note that only the fraction of the template within the mask is used for TM. The middle panels show the z-score maps obtained with the different masks, with mask heights indicated. Decreasing the height of the masks, i.e., the size of the template used, increases the background noise while decreasing the strength of the microtubule peaks.

**SI Fig. 8:**
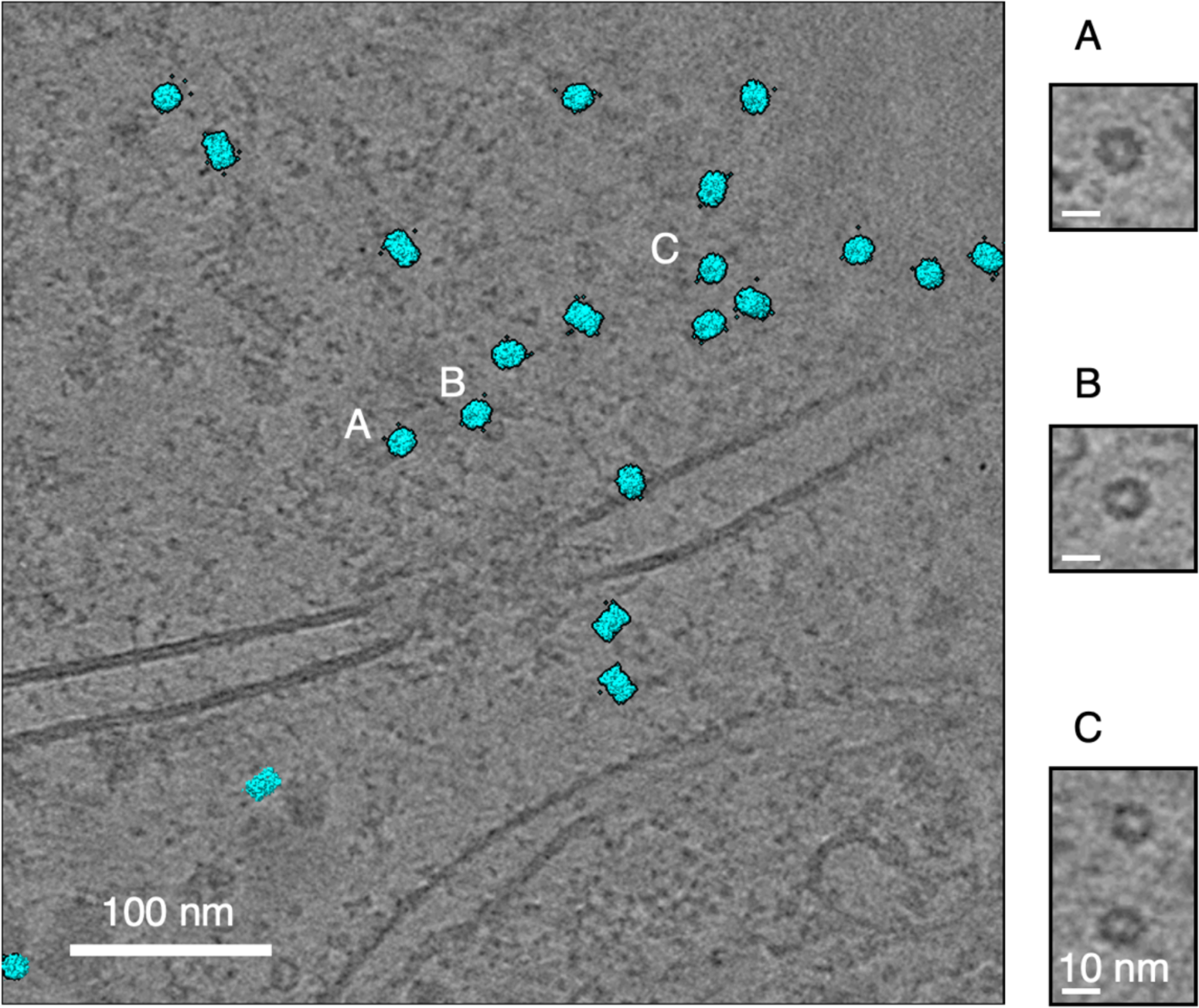
Template matching results for proteasome 20S. Templates (cyan) are repositioned to the position where TM reported high confidence peaks. A-C, Zoomed views of the tomogram at the positions identified by TM, showing that the peaks correspond to proteasomes.

